# Polyclonal HIV envelope-specific breast milk antibodies limit founder SHIV acquisition and cell-associated virus loads in infant rhesus monkeys

**DOI:** 10.1101/145524

**Authors:** Jonathon E. Himes, Ria Goswami, Riley J. Mangan, Amit Kumar, Thomas L. Jeffries, Joshua A. Eudailey, Holly Heimsath, Quang N. Nguyen, Justin Pollara, Celia LaBranche, Meng Chen, Nathan A. Vandergrift, James W. Peacock, Faith Schiro, Cecily Midkiff, Guido Ferrari, David C. Montefiori, Xavier Alvarez-Hernandez, Pyone Pyone Aye, Sallie R. Permar

## Abstract

Vertical HIV-1 transmission via breastfeeding is the predominant contributor to pediatric infections that are ongoing in this era of highly effective antiretroviral therapy (ART). Remarkably, only ~10% of infants chronically exposed to the virus via breastfeeding from untreated HIV-infected mothers become infected, suggesting the presence of naturally protective factors in breast milk. HIV-specific maternal antibodies are obvious candidates as potential contributors to this protection. This study assessed the protective capacity of common HIV envelope-specific non-broadly neutralizing antibodies isolated from breast milk of HIV-infected women in an infant rhesus monkey (RM), tier 2 SHIV oral challenge model. Prior to oral SHIV challenge, infant RMs were i.v. infused with either a single weakly-neutralizing monoclonal antibody (mAb), a tri-mAb cocktail with neutralizing and ADCC functionalities, or an anti-influenza HA control mAb. Of these groups, the fewest tri-mAb-treated infants developed plasma viremia (2/6, 3/6, and 6/8 animals viremic in tri-mAb, single-mAb, and control mAb groups, respectively). Tri-mAb-treated infants demonstrated significantly fewer transmitted/founder SHIV variants in plasma and decreased peripheral CD4+ T cell proviral loads at 8 week post-challenge compared to control mAb-treated infants. Abortive infection was observed as detectable CD4+ T cell provirus in non-viremic control mAb- and single-mAb-, but not tri-mAb-treated animals. Taken together, these results support the potential viability of maternal or infant vaccine strategies that elicit non-broadly neutralizing antibodies to prevent vertical transmission of HIV through breastfeeding.

## Importance

Due to the ongoing global incidence of pediatric HIV-1 infections even with wide antiretroviral therapy (ART) availability, many that occur via breastfeeding, development of vaccine strategies capable of preventing vertical HIV transmission through breastfeeding remains an important goal for achieving an HIV-free generation. Interestingly, in the absence of ART, only ~10% of infants chronically exposed to HIV orally via breastfeeding become infected, suggesting natural protective mechanisms in the remaining ~90% of HIV-exposed infants. In this study, we demonstrate that prophylactic infusion of non-broadly neutralizing human breast milk polyclonal antibodies with diverse anti-HIV functionalities limited acquisition and/or persistence of simian/human immunodeficiency virus (SHIV) following oral SHIV challenge of infant rhesus monkeys compared to controls. These findings imply that non-broadly neutralizing maternal antibodies may contribute to the inefficient vertical transmission observed in natural breastfeeding, and vaccine strategies eliciting such responses may prove viable for further limiting mother-to-child-transmission (MTCT) of HIV through breastfeeding.

## Introduction

According to the 2016 UNAIDS report, approximately 150,000 pediatric infections occur annually, accounting for ~10% of new global HIV-1 infections (1, 2). The benefits of breastfeeding to infant health are well recognized, yet vertical transmission of HIV-1 via breastfeeding results in nearly half of the annual mother-to-child-transmission (MTCT) occurrences (3). In resource-limited areas, formula-fed infants exhibit high mortality rates due to respiratory and diarrheal illness (4, 5) and thus, formula feeding is not a viable strategy to reduce pediatric HIV transmissions. While administration of antiretroviral therapy (ART) to HIV-1 infected, breastfeeding mothers greatly reduces MTCT rates to below 5% (6), socioeconomic barriers to ART availability and compliance (7, 8), as well as acute maternal infections make it unlikely that ART-based strategies alone can achieve eradication of pediatric HIV-1 (9–11). Therefore, developing effective immune-based prevention strategies, such as a maternal or infant vaccine to protect infants from oral HIV-1 acquisition during breastfeeding, may greatly contribute to the goal of achieving an HIV-free generation (12).

Interestingly, despite chronic mucosal HIV exposure multiple times daily during up to two years of breastfeeding, only ~10% of nursing infants of untreated HIV-infected mothers will acquire HIV (12), suggesting the presence of protective factors in breast milk. Thus, a better understanding of the naturally protective components of breast milk, such as mucosal antibodies, are of paramount importance in developing effective prophylactic vaccine strategies. Breast milk contains HIV-1 envelope (Env)-specific antibodies and Env-specific memory B cells (13, 14), both of which are primarily IgG1 isotype and are otherwise similar in specificity and function to those identified in blood of chronically infected individuals (15). Yet, the contribution of breast milk antibodies to the inefficiency of HIV-1 transmission through breastfeeding remains undefined. Induction or passive infusion of broadly neutralizing antibodies (bNAbs) is an attractive immunologic strategy for global HIV control (reviewed in (16)) including in the setting of postnatal HIV transmission (17, 18). Yet, bNAbs only develop naturally in fewer than 20% of individuals, typically take 2–4 years to develop after infection (19), and have been unable to be elicited through vaccination. Moreover, bNAbs have not been identified in breast milk (13, 20). Thus, it is unlikely that naturally acquired bNAbs largely contribute to the protection observed in the setting of MTCT through breastfeeding.

While achieving an HIV-free generation through vaccine development remains a priority, work to develop an effective treatment leading to remission or cure for infant breakthrough infections is also underway (21). One of the major goals of achieving HIV remission or cure is to reduce the size of the HIV reservoir to lengthen the time to viral rebound, and to make the host immune system competent to control residual virus autonomously (22). Currently, the major barrier for preventing HIV rebound after virologic control is the efficient ability of the virus to establish latency in resting memory CD4^+^ T cells (23, 24). Studies have demonstrated that this latent viral reservoir is seeded rapidly after SIV infection of rhesus monkeys, even before viremia is detectable in the plasma, and ART treatment as early as 3 days post-infection is too late to prevent establishment of the latent reservoir (25). Thus, the window for intervention before establishment of the viral reservoir is extremely limited and early ART alone may not be sufficient to attain sustained virologic remission. Post-exposure prophylaxis of single monoclonal broadly neutralizing antibodies (bNAbs) or a cocktail of potent bNAbs have been found to have promising effects in suppressing breakthrough infections and reducing the size of the viral reservoir in established infection (26–28). Important to the setting of infant HIV-1 infection, Hessell et al. have shown that administration of a cocktail of two potent neutralizing antibodies VRC07-523 and PGT121 at 24 hr post-exposure can reduce the establishment of permanent viral reservoirs in infant rhesus monkeys (18). Hence, potent neutralizing antibodies could contribute to reducing the size of viral reservoirs early after establishment, and might be a synergistic strategy with ART to obtain complete virologic control and delay plasma viral rebound.

Infants orally infected during breastfeeding acquire antibodies present in breast milk concurrently with the virus. Yet, the in vivo contributions of breast milk-derived antibodies to the inefficiency of breast milk HIV-1 transmission and viral reservoir establishment remain largely unexplored. In this study, we sought to define the impact of passively infused and orally dosed breast milk-derived monoclonal antibodies (mAbs) on infant oral virus acquisition and dissemination in the periphery and lymphoid tissues. MAbs selected for this study were isolated from breast milk B cells of a cohort of HIV-1-infected Malawaian women and were intended to represent breast milk IgG antibodies with various antiviral functionalities and envelope specificities (20). RMs were prophylactically passively infused with the maternal breast milk mAbs to mimic antibody transfer via the placenta, and then orally-exposed to these breast milk-derived mAbs during repeated oral low dose challenge with the tier 2 chimeric simian/human immunodeficiency virus, SHIV-1157ipd3N4 (29). Defining the contributions of non-broadly neutralizing breast milk-derived antibodies to the naturally inefficient transmission of HIV-1 through breastfeeding may inform the design of maternal and infant vaccines aimed at eliminating postnatal HIV-1 infections and limiting the size of the viral reservoir in the setting of breakthrough infections.

## Results

### Selection of maternal breast milk mAbs for in vivo evaluation in infant monkeys and study design

The HIV Env-specific mAbs isolated from breast milk B cells of lactating, HIV-1-infected Malawian women (14) and selected for infusion into infant RMs in this study were initially characterized based on binding specificity, epithelial and dendritic cell-virus binding inhibition, ADCC, and neutralization against the tier 2 challenge virus in this study, SHIV-1157ipd3N4 (29), as well as neutralization of several tier 1 HIV/SHIV variants (**Figure 1A**) (20). As previously reported, all of the mAbs isolated from milk B cells were IgG1, did not demonstrate broadly neutralizing activity, and had similar characteristics to those isolated from peripheral blood HIV-1 Env-specific B cells (14). The infusion mAbs were selected based on their diversity of functions and specificities; mAb DH378 demonstrated CD4-blocking capability associated with CD4 binding site (CD4bs)-specificity and mAb DH377 demonstrated linear and conformational variable loop 3 (V3) binding, both of which are specificities previously associated with decreased MTCT risk (30, 31), and mAb DH382 demonstrated constant region 1 (C1) specificity and competed with mAb A32-Env binding. All of these anti-HIV mAbs were able to block epithelial and dendritic cell-virus binding in vitro. Unsurprisingly, the A32-like mAb DH382 was capable of potent ADCC activity against the tier 2 challenge SHIV-1157ipd3N4, with a maximum % specific killing of 40.7% and an endpoint concentration of <0.04µg/mL. Both DH378 and DH377 demonstrated neutralization potency against all tier 1 viruses tested, including SHIV SF162P4, SHIV BaL-P4, MW965.26, and SHIV-1157ipEL-p. Of the mAbs isolated from maternal milk, only DH378 demonstrated weak neutralization activity against the tier 2 clade C SHIV challenge virus (IC_50_=79µg/mL).

**Figure 1.**
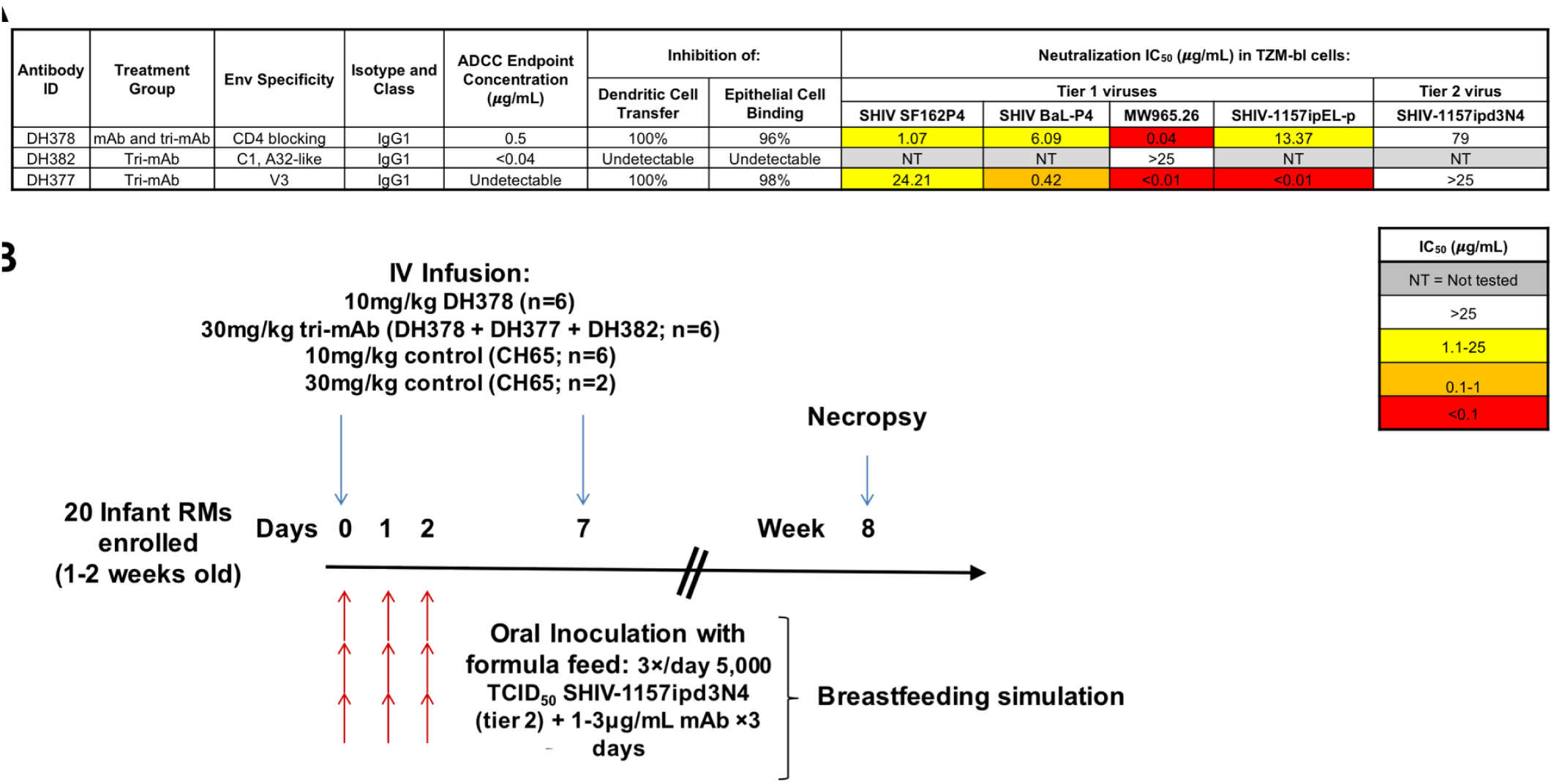
Infant RM passive mAb infusion and oral SHIV challenge study design. **A)** Functional characterization of maternal breast milk mAbs selected for passive infusion into infant RMs. MAbs were selected based on diverse functionality, including Env binding specificity and ADCC, inhibition of dendritic cell virus transfer, inhibition of epithelial cell-virus binding, and virus neutralization functionalities. **B)** Timeline for passive mAb infusion and SHIV-1157ipd3N4 challenge of infant RMs. Twenty infant RMs (1–2 weeks old) were systemically infused with the indicated mAbs and subsequently orally challenged after 1 hour with 5,000 TCID_50_ of the tier 2 SHIV-1157ipd3N4 in formula feed to mimic infant oral viral exposure during breast feeding. Animals were reinfused 7 days after the first mAb infusion with the same mAb dose to sustain systemic Ab titers. Animals were necropsied at 8 weeks after challenge and lymphatic and GI tract tissues were collected.

To assess the degree to which common HIV Env-specific breast milk antibodies contribute to protection against oral virus acquisition and replication, 20 infant RMs were divided into 2 anti-HIV-1 Env-specific breast milk mAb treatment groups and a control mAb treatment group for passive mAb infusion and oral SHIV-1157ipd3N4 challenge (**Figure 1B**). In this study scheme, systemic passive infusion was intended to simulate placental maternal IgG transfer, while the oral inoculum was intended to represent oral exposure to the combination of maternal breast milk HIV Env-specific antibodies and infectious virus. The first treatment group consisted of 6 animals infused with 10mg/kg of DH378 as this mAb exhibited detectable, albeit weak neutralization potency against the tier 2 challenge SHIV, as well as dendritic and epithelial cell-virus binding inhibition. The second group consisted of 6 animals infused with 30mg/kg of an equimolar mixture of mAbs DH378, DH377, and DH382—a mixture containing mAbs that demonstrate potent ADCC functionality, a variety of binding specificities including a specificity previously associated with decreased MTCT (V3) (30, 31), dendritic and epithelial cell-virus binding inhibition, and weak tier 2 neutralization against the challenge SHIV. Anti-influenza HA mAb CH65 was employed as a control mAb treatment in 8 animals and was infused at a dose of either 10mg/kg (n=6) or 30mg/kg (n=2) to provide an appropriate dose control for each anti-HIV mAb infusion group. Of note, control animals from both dose groups were combined for statistical comparisons. To simulate placental maternal Ab transfer, infants were infused intravenously with the mAbs 1 hour prior to the first of 9 oral SHIV-1157ipd3N4 challenges (3 times/day for 3 consecutive days). Each oral challenge consisted of 5,000 TCID_50_ SHIV-1157ipd3N4 incubated for 15 minutes on ice with 1µg/mL of the single mAb, or 3µg/mL of the tri-mAb mixture matching each infusion group, concentrations selected to mimic the levels of HIV Env-specific antibodies in breast milk (13). The SHIV/mAb mixture was diluted in 10mL of infant formula prior to infant feeding to simulate breastfeeding. Animals were reinfused with mAbs matching the initial infusion at 7 days to sustain systemic mAb titers. SHIV-challenged infant monkeys were then followed for plasma mAb kinetics, systemic cell-free and cell-associated virus loads, and were necropsied at 8 weeks for assessment of lymphoid and GI tract tissue viral loads (**Figure 1B**).

### Pharmacokinetics of maternal milk HIV-1 mAb infusion of infant RM serum and saliva

Serum mAb concentrations at longitudinal time points through 8 weeks post SHIV-challenge were measured by ELISA to determine the kinetics of the mAb persistence. Serum mAb levels in animals from all treatment groups were largely similar through 2 weeks post-infusion achieving concentrations of 9.2×10^4^–4.5×10^6^ ng/mL, but some variability arose thereafter as certain animals cleared the infused mAbs more rapidly than others (**Figure 2A**; **Figure S1**). Additionally, the natural SHIV Env-specific antibody responses likely developed in a subset of DH378 and tri-mAb treated infants after 28 days, as the Env-binding IgG response did not decline to baseline in these infants.

**Figure 2.**
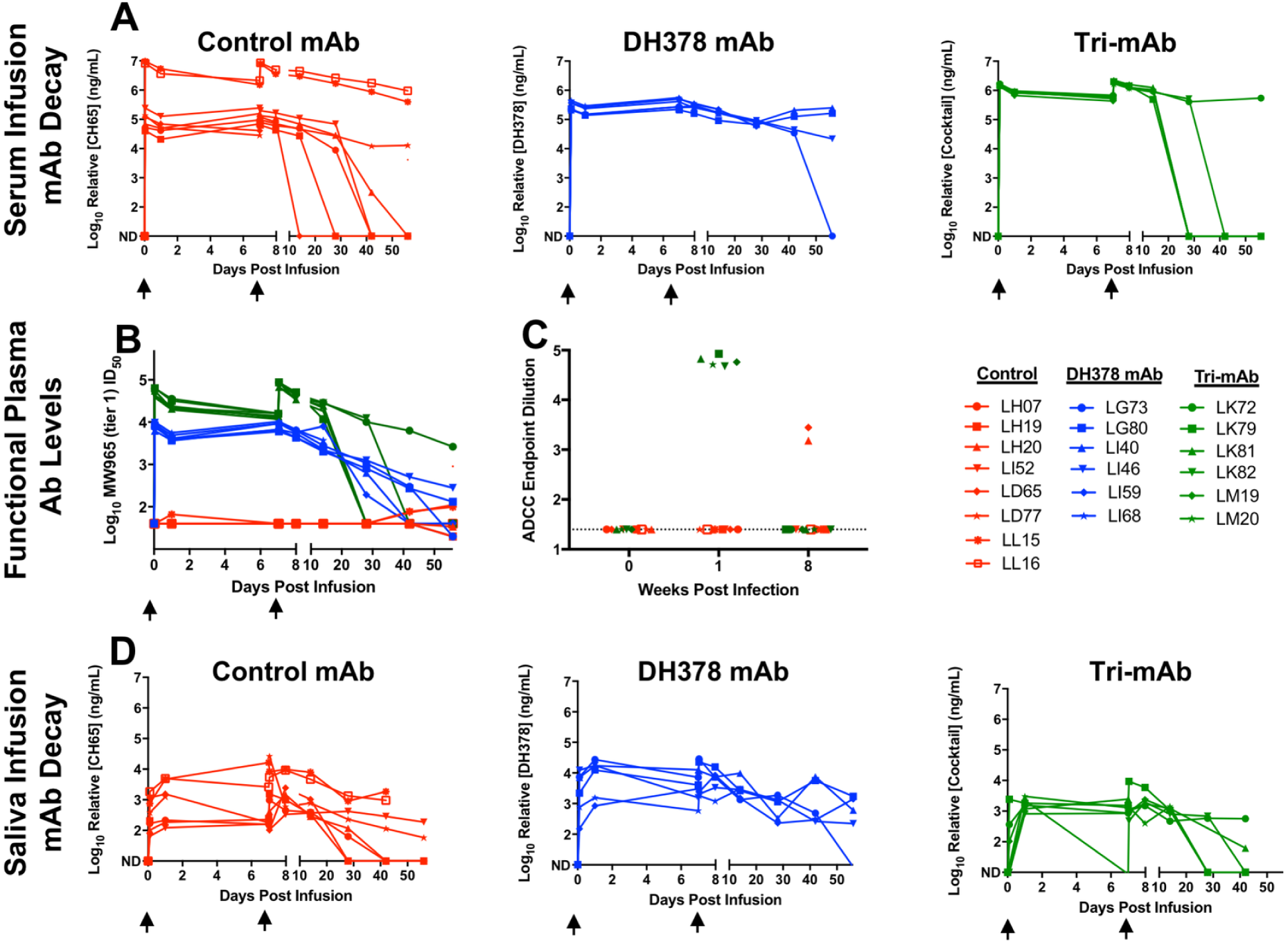
The kinetics of passively infused mAbs and anti-SHIV Ab function in passively breast milk mAb-infused, SHIV-challenged infant RM serum and saliva. Concentrations of infused mAbs in **A)** serum and **D)** saliva from pre-infusion to 8 weeks post infusion are depicted for control mAb-treated (CH65; red), DH378 mAb-treated (DH378; blue), and tri-mAb-treated (DH378, DH377, DH382; green) infant RMs. **B)** Neutralizing Ab levels in serum reported as ID_50_ against the tier-1 virus MW965 from pre-infusion to 8 weeks post challenge. **C)** ADCC-mediating Ab levels in serum reported as endpoint titer against SHIV-1157ipd3N4 at preinfusion, 1 week (after 2^nd^ mAb infusion), and 8 weeks post challenge. Black arrows indicate systemic mAb infusions at days 0 and 7. ND indicates not detectable.

Plasma neutralizing antibody (NAb) and ADCC levels were measured in all infused infant RMs to assess in vivo functionality of the infused treatment mAbs and development of host humoral responses (**Figure 2B, C**). Predictably, NAb responses against the tier 1 clade C virus MW965 were high in all DH378- and tri-mAb-treated animals early after mAb infusion, and the magnitude of the responses declined to undetectable in some animals as early as 6 weeks post challenge as circulating mAb levels deteriorated (**Figure 2B**). Interestingly, several control animals began to develop a natural tier 1, MW965 NAb responses as early as 6 weeks post-challenge. None of the animals in this study exhibited neutralization against either the tier 2 SHIV-1157ipd3N4 challenge virus or the related tier 1 virus SHIV-1157ipEL-p (32) at peak mAb concentrations or 8 weeks post challenge, which was unsurprising as serum mAb concentrations never surpassed that of the in vitro IC_50_ (**Figure 1A**). Serum ADCC endpoint titers against SHIV-1157ipd3N4-infected CD4+ T cells were measured preinfusion, at peak infusion mAb concentration (1 hour after second infusion), and at 8 weeks post initial infusion for control and tri-mAb-treated RMs (**Figure 2C**; DH378-treated RMs not tested). As expected, serum from tri-mAb-treated animals exhibited detectable ADCC function, which deteriorated by 8 weeks post initial infusion. Interestingly, 2/8 control mAb-treated animals, but none of the tri-mAb-infused infant RMs, developed detectable autologous ADCC-mediating mAbs against the challenge SHIV strain after infection (8 weeks post initial infusion).

Saliva mAb concentrations were also measured prior to and following oral challenge (**Figure 2D**). All animals demonstrated detectable mAb levels in saliva at 1 hour post infusion, which was the time of the first SHIV challenge. Saliva mAb levels peaked around 1 day post infusion in all animals and only slightly declined by 7 days post initial infusion in most animals. The tri-mAb-treated animal, LK79, was unique in that it cleared the infused mAb rapidly from the saliva, demonstrating undetectable mAb treatment levels in saliva by 7 days post initial infusion, as well as clearance of the infused mAbs after the second infusion by day 28 of the study.

### Effect of maternal milk HIV-1 antibody treatment on SHIV acquisition and plasma viremia in orally challenged infant RMs

To characterize the effectiveness of the mAb treatments in preventing SHIV acquisition, SHIV plasma viral RNA loads of orally-challenged infant RMs were measured in longitudinal samples by qRT-PCR. 6/8 (75%) CH65-treated control animals, 3/6 (50%) DH378-treated animals, and 2/6 (33%) tri-mAb cocktail-treated animals became detectably viremic after 9 oral SHIV challenges resulting in statistically similar SHIV acquisition rates between controls and each treatment group (**Figure 3A**; FDR corrected Fisher’s exact test; Controls vs. DH378 p=0.58, tri-mAb cocktail p=0.55). Peak and set point viral RNA loads in plasma were largely similar between CH65-treated control animals (median and range; peak=6.2×10^6^copies/mL, 9.3×10^5^–2.7×10^7^copies/mL; set point=4.4×10^5^copies/mL, 5.4×10^4^–9.5×10^7^copies/mL) and tri-mAb-treated animals (median and range; peak=2.1×10^6^copies/mL, 5.3×10^5^–3.6×10^6^copies/mL; set point=3.0×10^5^copies/mL, 1.2×10^4^–5.9×10^5^copies/mL) when excluding animals with undetectable viral RNA loads (**Figure 3B**). Although, the tri-mAb cocktail-treated animal LK81 emerged as an outlier with the lowest plasma peak and set point viral RNA loads observed in a viremic animal from this study (peak=5.3×10^5^copies/mL; set point=1.2×10^4^copies/mL). Interestingly, while set point viral RNA loads in plasma were similar between viremic CH65-treated control animals and viremic DH378-treated animals (median and range; peak=7.7×10^7^copies/mL, 7.4×10^7^–1.1×10^8^copies/mL; set point=5.1×10^5^copies/mL, 3.5×10^5^–8.9×10^6^copies/mL), the peak viral RNA loads in plasma from viremic DH378-treated animals was significantly higher than that from viremic controls (**Figure 3B**; FDR-corrected Wilcoxon test; FDR-corrected p=0.05). Of note, DH378-treated animal LI68 died due to unrelated causes (choking event) prior to the 4-week time point, resulting in insufficient sample for proviral load quantification. Yet, its 2-week viral RNA load was undetectable and no other viremic animal in the study demonstrated an initial detectable viral RNA load after 2 weeks, thus, we considered LI68 uninfected. Given the ranges of peak and set point viral RNA loads, MHC typing was conducted on all animals. Several animals possessed MHC alleles highly (Mamu-A001, Mamu-B017) or weakly (Mamu-A002, Mamu-B047) associated with low set point viral loads and/or longer survival lengths (**Table S1**) (33–40). However, these protective alleles were not more common in any single treatment group (67% for control mAb-, and tri-mAb-treated animals, 80% for DH378 mAb-treated animals). Additionally, all groups also contained animals with the Mamu-A004 allele, which is associated with increased set point viral loads (41). Interestingly, all viremic, DH378 mAb-treated animals possessed this allele, which could contribute to the significantly higher plasma viral loads observed in these animals (**Figure 3B**).

**Figure 3.**
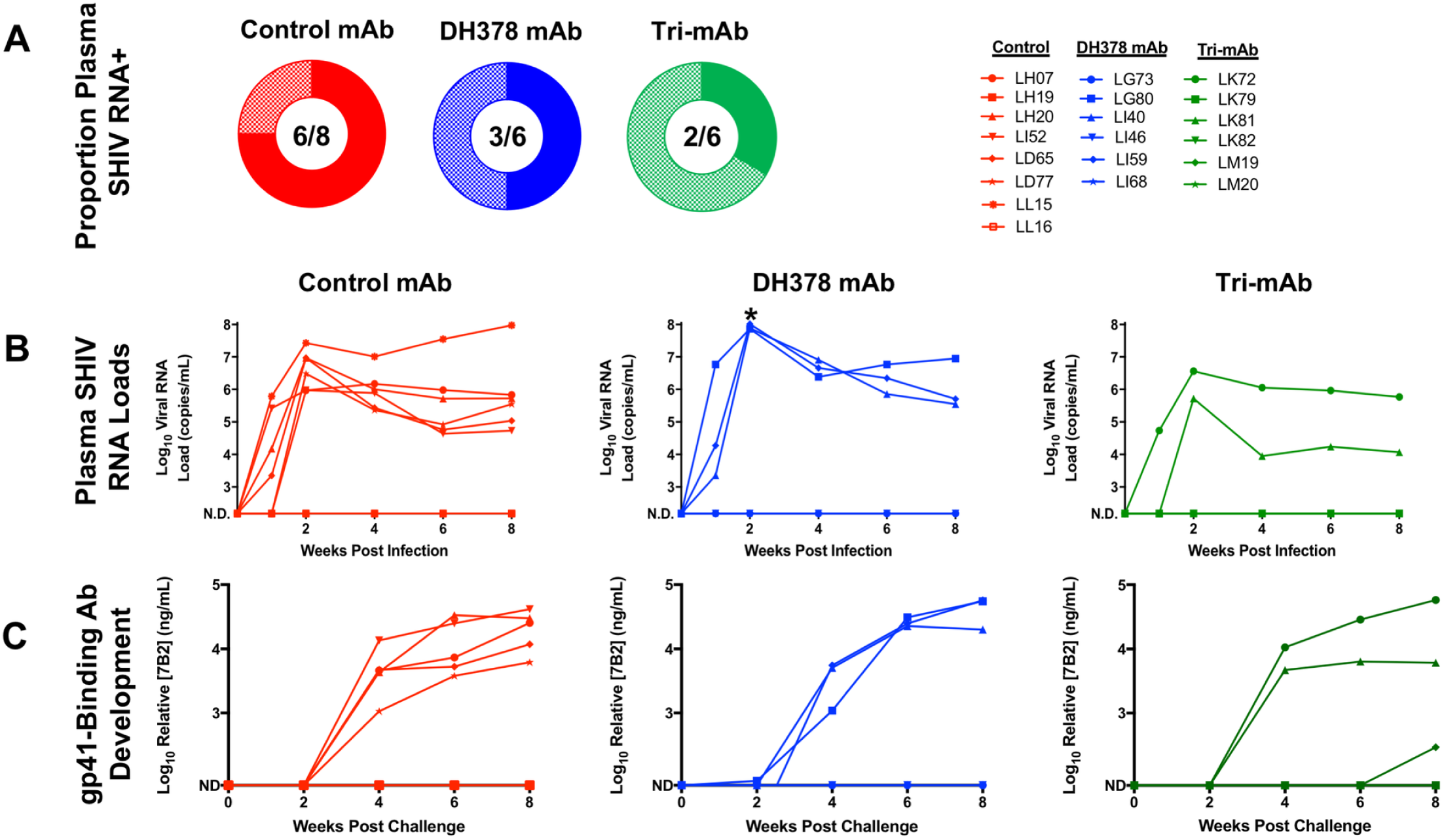
Plasma viral loads, infection rates, and Env gp41 seroconversion of passively breast milk mAb-infused, SHIV-challenged infant RMs. **A)** Proportion of viremic RMs with 6/8 control mAb-treated RMs (red), 3/6 DH378 mAb-treated RMs (blue; Fisher’s exact test; FDR-corrected p=0.58), and 2/6 tri-mAb-treated RMs (green; Fisher’s exact test; FDR-corrected p=0.55) demonstrating viremia as defined by a detectable plasma viral RNA load at any tested time point. **B)** Plasma viral RNA loads in mAb treated, orally SHIV challenged infant RMs. * indicates significantly higher 2 week post infection plasma viral RNA loads in DH378 mAb-treated RMs compared to CH65-treated RMs (Wilcoxon test; FDR-corrected p=0.05). **C)** Env gp41-binding serum Ab responses (measured as 7B2 Ab mEq) from preinfusion to 8 weeks post-infusion in infant RMs. ND indicates not detectable.

To assess the development of virus-specific antibodies and seroconversion after infection, anti-HIV gp41 antibody titers in serum were measured against HIV Env gp41. As the infusion mAbs were gp120-specific, detection of gp41-specific binding in longitudinal serum represented the host antibody response. All but one animal with detectable plasma viral RNA developed strong anti-gp41 antibody responses by 4 weeks post-challenge (**Figure 3C**). Control mAb-treated animal LL15, who failed to develop a gp41-directed antibody response, sustained the highest week 8 viral loads and the greatest PBMC CD4+ T cell depletion observed in this study (**Figure S2**). Interestingly, only 1 tri-mAb-treated animal lacking detectable viremia, LM19, developed a detectable Env gp41-specific antibody response at week 8 post-challenge, but this response was ~2 logs lower in magnitude than those of the viremic animals and detected at only the 8 week necropsy time point, suggesting that this may be a cross reactive response.

### Cell-associated SHIV load in blood and tissues from breast milk mAb-treated, orally SHIV-challenged infant RMs

To determine whether the presence of pre-existing HIV-1 Env-specific mAbs impacted the cell-associated viral load in blood and tissue compartments following oral challenge in mAb-infused infant RMs, multiple approaches were employed to quantitate SHIV virus load in PBMCs, GI tract, and lymphoid tissues of SHIV-challenged infant RMs. These approaches included quantifying tissue mononuclear cell proviral loads, proportions of viral RNA producing T cells, and tissue-associated infectious virus titers (**Figure 4**). SHIV provirus in CD4+ T cells isolated from PBMCs (preinfection, 2 and 8 weeks post-infection), GI, and lymphoid tissues at 8 weeks post challenge and measured via ddPCR in viremic infants demonstrated widely variable, but routinely detectable proviral loads (10^4^–10^7^ gag copies/million CD4+ T cells) independent of tissue type or mAb treatment (**Figure 4A**). Interestingly, SHIV provirus was undetectable in PBMCs or any tissues isolated from tri-mAb-treated animal LK81, which exhibited detectable, albeit relatively low plasma SHIV RNA loads.

**Figure 4.**
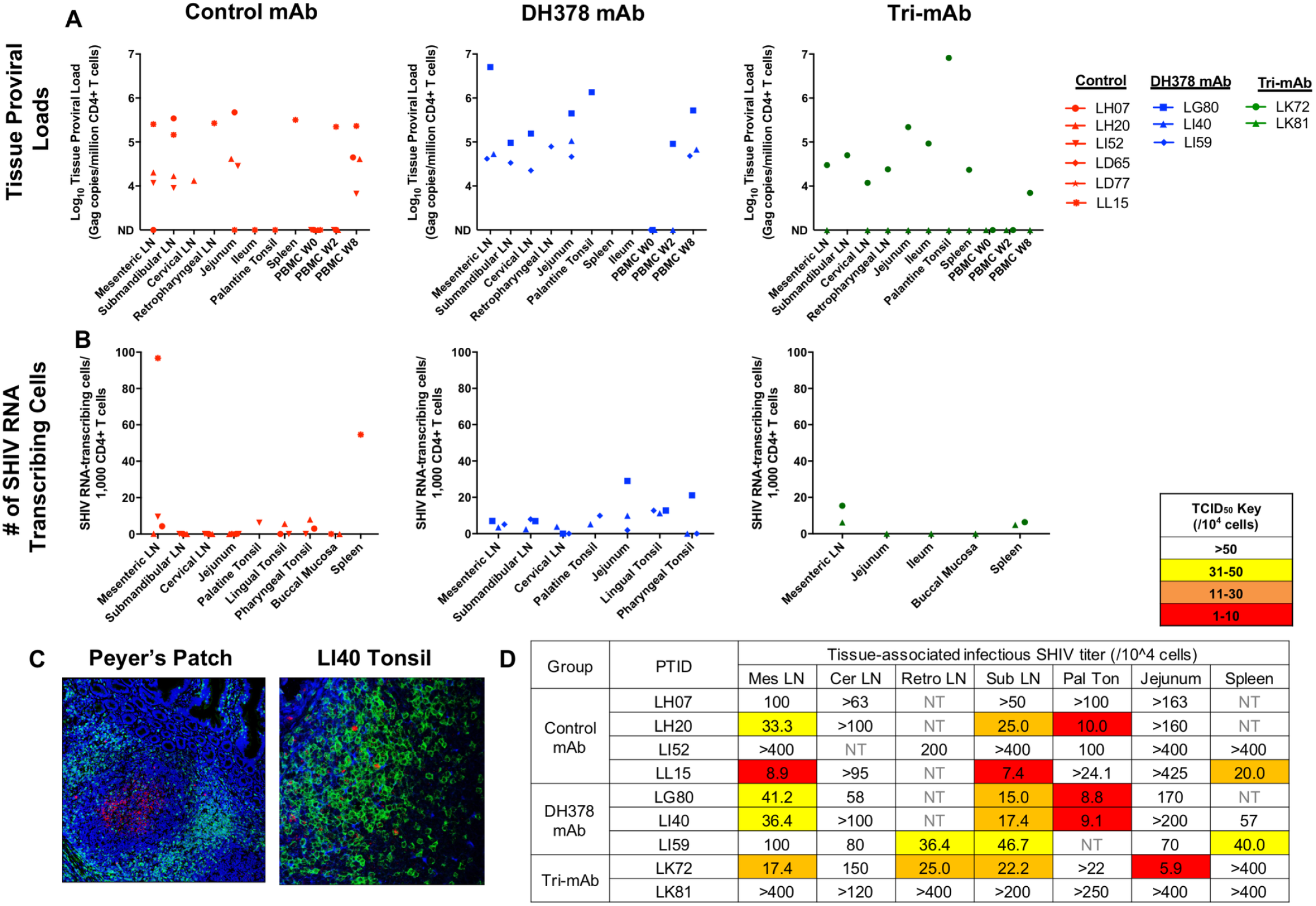
Tissue-associated SHIV measures in mononuclear cells isolated from lymphoid and GI tissues of viremic infant RMs 8 weeks post passive breast milk mAb infusion and oral SHIV challenge. **A)** CD4+ T cell proviral DNA loads reported as copy number/million CD4+ T cells in lymphoid and GI tissue mononuclear cells and **B)** number of SHIV RNA producing CD3+ T cells in lymphoid and GI tissue blocks reported as the number of SHIV Gag RNA+ cells per 1,000 CD3+ T cell were measured for control mAb-treated (red), DH378 mAb-treated (blue), and tri-mAb-treated RMs (green) through ddPCR and in situ hybridization, respectively. ND indicates not detectable. **C)** Representative images of stained tissue blocks demonstrating active SHIV RNA transcription within a Peyer’s Patch (LI40 jejunum) and in CD3+ T cells in lymphoid tissue (LI40 lingual tonsil) employed to assess the number of SHIV RNA transcribing cells within lymphoid and GI tissues. Blue, green, and red depict nuclear, CD3, and SHIV Gag RNA staining, respectively. **D)** Tissue-associated infectious SHIV-1157ipd3N4 titers measured through tissue mononuclear cell coculture with TZM-bl reporter cells. Reported titers represent the estimated minimum number of mononuclear cells required to yield detectable infection (cutoff defined by mononuclear cells from unchallenged RM negative controls) of the TZM-bl cells in 50% of replicates. The cutoff for The limit of detection varied for each sample and was determined by mononuclear cell availability. NT indicates untested samples.

The prevalence of viral RNA expressing cells in tissues was measured by in situ hybridization assays and quantified as the number of viral RNA producing cells per 1,000 CD3+T cells (**Figure 4B-C**). The proportion of SHIV RNA transcribing cells was largely similar in lymphoid and GI tissues from animals in all treatment groups. However, mesenteric lymph node and spleen from control mAb-treated animal LL15 exhibited a notably higher prevelance of SHIV RNA positive cells compared to other SHIV-infected infants (**Figure 4B**). Additionally, extensive SHIV RNA production was observed within, and to a lesser degree surrounding, Peyer’s patches of the small intestine as well as in CD3+ T cells of lymphoid tissues (**Figure 4C**). Interestingly, SHIV RNA detection within the Peyer’s patches largely failed to colocalize with CD3 expression, indicating that virus associated with dendritic cells and other APCs within these germinal centers contribute to this SHIV RNA detection.

To assess the level of cell-associated infectious SHIV within oral-associated lymphoid and GI tissues at 8 weeks post-challenge, tissue mononuclear cells were isolated, serially diluted, and cocultured with TZM-bl reporter cells with luminescence output indicating the degree of infectious SHIV production (**Figure S3**). Tissue-associated infectious virus titer was calculated as the number of cells required to sustain detectable infection in 50% of the replicates, with the detection threshold established as 3 standard deviations above the mean luminescence output of PBMCs from 3 naive RMs (3,019 RLU). Of note, cell number availability was highly variable between samples, resulting in a range of detection limits (range=2×10^5^–4×10^6^ mononuclear cells) and number of replicates (2–8 replicates) for the assay. In general, tissue-associated infectious SHIV titers were similar between treatment groups (**Figure 4D**). However, within each animal the palatine tonsil exhibited the highest cell-associated infectious titers of any tissue in 4 of 7 infants with sufficient tonsil tissue for the assay. As this assay employed total mononuclear cells instead of isolated CD4+ T cells, this observation could be attributed to variability in CD4+ T cell proportion of total mononuclear cells between tonsils and other lymphoid tissues. However, flow cytometric analysis revealed that proportions of CD4+ T cells of total CD45+ mononuclear cells were lower in palatine tonsil than other lymphoid tissues (**Figure S4**), indicating a potential highly infectious tissue-associated virus titer in palatine tonsil. Interestingly, two viremic animals, CH65-treated animal LH07 and tri-mAb-treated animal LK81, exhibited undetectable tissue-associated infectious virus titers. Yet, the tissue shipment for LH07 experienced unexpected delays resulting in prolonged incubation of tissues in culture media prior to cell isolation, which may have led to inactivation of SHIV-producing cells, whereas LK81 had the lowest plasma viral load of all viremic animals and undetectable blood cell-associated virus load.

Proviral loads, tissue-associated infectious virus titers, and tissue viral RNA levels in PBMCs (preinfection, 2 and 8 weeks post-infection), GI tract tissues, and lymphoid tissues in non-viremic, SHIV-challenged infant RMs were also measured to assess the possibilities of abortive infection or undetectable, low level replication in the non-viremic infant RMs (**Figure 5**). Interestingly, several animals with undetectable plasma SHIV RNA loads throughout the study including DH378-treated animals LG73 and LI46, and CH65-treated animal LH19, exhibited detectable SHIV proviral DNA loads in various tissues at similar levels to those measured from animals with detectable plasma SHIV RNA loads (**Figure 5A**). To confirm the presence of tissue-associated virus in these non-viremic animals, the SHIV proviral envelopes were amplified by PCR of genomic DNA (gDNA) from submandibular LN CD4+ T cells of LH19 and LG73 and sequenced to demonstrate the presence of full-length envelope open reading frames (**Figure 5B**). One SHIV *env* variant from each animal was cloned and cotransfected in 293T cells with the SG3Δ*env* backbone to generate pseudovirues in order to assess *env* functionality of the provirus. Pseudovirus infectious functionality was measured as a tat-regulated increase in relative luminescence units (RLUs) after incubation with TZM-bl reporter cells. Both pseudoviruses were capable of infecting TZM-bl reporter cells with the LH19 and LG73 *env* pseudoviruses eliciting luminescence magnitudes of 571,294 RLU and 176,820 RLU, respectively, which was appreciably higher than that of cell only controls (~500 RLU).

**Figure 5.**
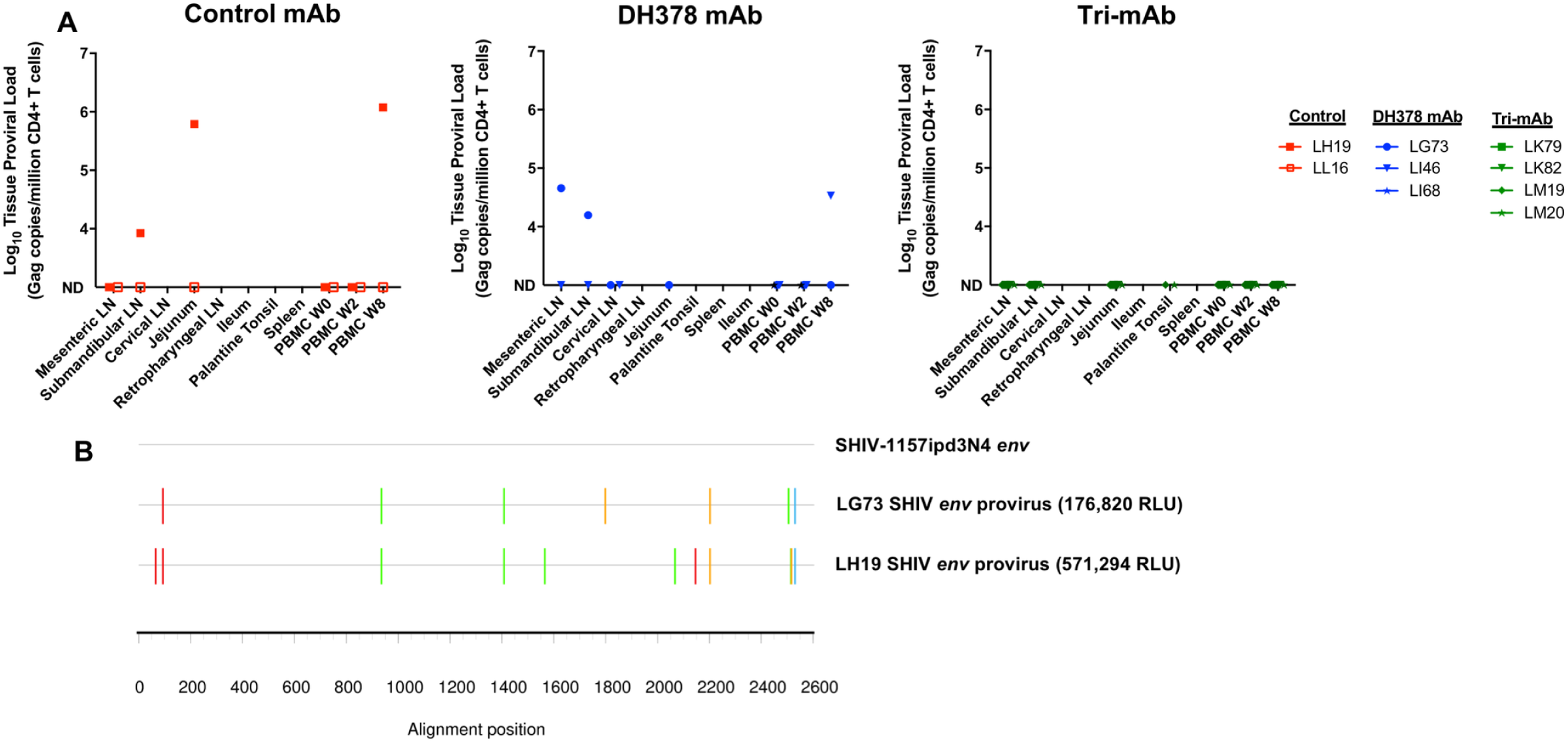
Tissue-associated SHIV DNA in CD4+ T cells isolated from lymphoid and GI tissues of non-viremic infant RMs 8 weeks post passive breast milk mAb infusion and oral SHIV challenge. **A)** Proviral DNA loads reported as copy number/million CD4+ T cells in lymphoid and GI tissue mononuclear cells were measured for control mAb-treated (red), DH378 mAb-treated (blue), and tri-mAb-treated RMs (green) through ddPCR. ND indicates not detectable. **B)** Highlighter plot alignments of 2 amplified SHIV *env* sequences obtained through SGA of extracted DNA from CD4+ T cells isolated from LH19 and LG73 submandibular lymph nodes compared to the SHIV-1157ipd3N4 *env* stock sequence. Colored lines indicate nucleotide mismatches compared to the SHIV stock. These *envs* were cloned and subsequently incorporated into pseudoviruses with a SG3∆*env* backbone through co-transfection in 293T cells. Pseudovirus infectivity was estimated in TZM-bl reporter cells as RLUs compared to cell-only controls (<500 RLU). Pseudoviruses expressing *env* isolated from LH19 and LG73 demontstrated infectious capability with 571,294 RLU and 176,820 RLU magnitudes, respectively.

Thus, 4 infants, at least 1 from each mAb treatment group, had discordance between the presence of detectable plasma viral RNA and CD4+ T cell provirus (**Figure S5A**). Provirus was observed in CD4+ T cells isolated from non-viremic animals LG73, LI46, and LH19. Alternatively, provirus was not detectable in the viremic animal LK81. Interestingly, prior to correction for multiple comparisons, tri-mAb-treated infants, but not DH378-treated infants, demonstrated lower magnitude proviral loads in 8 week PBMCs compared to control mAb-treated animals (Wilcoxon test; raw p=0.03; FDR-corrected p=0.18). Taken together, 7/8 CH65-treated control animals, 5/6 DH378-treated animals, and 2/6 tri-mAb cocktail-treated animals demonstrated either detectable viremia or proviral loads following oral SHIV challenge (**Figure S5B**).

To assess whether breast milk HIV Env-specific mAb pretreatment had an impact on the CD4+ T cell dysfunction in infected infants, a transcriptome profile of CD4+ T cells isolated from provirus containing lymphoid and GI tissues of infant RMs was also assessed. The genes included in the analysis were selected for their known involvement in inflammatory processes, particularly CD4+ T cell activation and exhaustion, and cell death pathways. Clustering analysis of relative gene RNA expression in CD4+ T cells from various infant RMs and tissues revealed clustering independent of tissue type, and mAb treatment group, indicating that infection in the presence of preexisting HIV Env-specific mAb did not affect target cell profiles in infected infants (**Figure S6**). Tissue and PBMC CD4+ T cell gene transcription profiles did cluster loosely corresponding to CD4+ T cell proviral loads.

### Plasma SHIV variant diversity in mAb-treated and orally SHIV-infected infant RMs

To assess the potential impact of maternal breast milk mAb passive infusion of infants on the genetic bottleneck of mucosal virus transmission and SHIV *env* diversification by peak viremia (2 weeks post challenge), single SHIV variants were isolated from infant RM plasma using single genome amplification (SGA). Of note, the median viral diversity of the SHIV-1157ipd3N4 challenge virus stock was 0.02% (range=0–0.07%), appreciably lower than the diversity typically observed in chronically HIV-1-infected humans (>1% diversity). Viral diversity within each animal was visualized with phylogenetic tree and highlighter plot (**Figure S7**). Distinct SHIV transmitted/founder (T/F) variants present in plasma isolated at peak viremia (2 weeks post challenge) were enumerated using previously established criteria (42), which required that a genotypically distinct variant contain 2 novel mutations (to minimize the possibility of recombination yielding false results), and that these mutations be observed in at least 2 SGAs (to minimize the effects of PCR-related mutation). This process demonstrated a similar number of distinct SHIV variants in plasma at peak viremia between control mAb-treated RMs (**Table 1**; n=8; median variants isolated=2.5; range=0–4) and DH378 mAb-treated animals (n=6; median variants isolated=0.5; range=0–2; Wilcoxon test; raw p=0.16; FDR-corrected p=0.26). However, prior to correction for multiple comparisons, plasma SHIV diversity at peak viremia was lower in tri-mAb-treated RMs (n=6; median variants isolated=0; range=0–1; Wilcoxon test; raw p=0.05; FDR-corrected p=0.18), indicating a possible sieve effect of the tri-mAb treatment on oral SHIV acquisition. When considering SHIV *env* amplicons from all RMs in a single tree rooted to the SHIV-1157ipd3N4 challenge virus *env*, the tri-mAb-treated RMs LK72 and LK81 are more homogenous compared to the other 2 infusion groups (**Figure S8**).

**Table 1.** Diversity of SHIV-1157ipd3N4 variants in serum isolated 1 week post initial oral challenge from control mAb-, DH378 mAb-, or tri-mAb-treated infant RMs.

| Control mAb<br>(Animal ID) | # of<br>Variants | SGA<br>Sequences per<br>Animal | DH378 mAb<br>(Animal ID) | # of<br>Variants | SGA<br>Sequences per<br>Animal | Tri-mAb<br>(Animal ID) | # of<br>Variants | SGA<br>Sequences per<br>Animal |
| --- | --- | --- | --- | --- | --- | --- | --- | --- |
| LI52 | 4 | 33 | LG80 | 2 | 30 | LK72 | 1 | 25 |
| LD65 | 3 | 36 | LI59 | 2 | 18 | LK81 | 1 | 32 |
| LH20 | 3 | 23 | LI40 | 1 | 21 | LK79 | 0 | - |
| LL15 | 3 | 27 | LI46 | 0 | - | LK82 | 0 | - |
| LH07 | 2 | 16 | LG73 | 0 | - | LM19 | 0 | - |
| LD77 | 1 | 42 | LI68 | 0 | - | LM20 | 0 | - |
| LH19 | 0 | - |  |  |  |  |  |  |
| LL16 | 0 | - |  |  |  |  |  |  |
| Median (Range)<br># Variants | 2.5 (0-4) |  | Median (Range)<br># Variants | 0.5 (0-2) |  | Median (Range)<br># Variants | 0 (0-1) |  |

## Discussion

Despite multiple times daily chronic oral exposure to HIV for up to 2 years, only ~10% of breastfeeding infants of untreated HIV-infected mothers acquire HIV (12). Furthermore, postnatally HIV-infected infants demonstrate slower disease progression compared to in utero or peripartum infected infants (43). Therefore, a better understanding of the naturally protective components in breast milk of HIV-infected mothers, including breast milk-derived antibodies, could aid in the development of effective vaccination strategies aimed at reducing MTCT and other modes of HIV acquisition. Passively infused bNAbs have been shown to protect orally challenged infant RMs from virus acquisition when provided prophylactically (17) or even 1 day post challenge (18). Yet bNAbs only develop naturally in a subset of infected individuals years after infection and have yet to be elicited by vaccination. The protective effects of maternal breast milk mAbs that lack neutralization breadth remains uncertain. Thus, we sought to probe the protective capabilities of several non-broadly neutralizing breast milk mAbs previously isolated from a cohort of HIV-infected Malawian women in an infant RM passive infusion and oral SHIV challenge model. The three maternal breast milk mAbs selected for inclusion in this study were intended to represent mAbs exhibiting various anti-HIV functionalities (ADCC, tier 1 and weak tier 2 neutralization, dendritic cell-virus binding inhibition, epithelial cell-virus binding inhibition) and HIV Env specificities (C1, V3, CD4-blocking). The treatment groups in this study were designed to assess whether a CD4-blocking mAb with tier 1 and weak autologous tier 2 neutralization could contribute to protection (DH378 mAb treatment group), or if polyclonal anti-viral functionalities also played an important role (tri-mAb treatment group).

The study scheme sought to model infant exposure to oral virus as would occur through breastfeeding via an HIV-infected mother. Therefore, we orally challenged animals 3 times daily for 3 days with SHIV premixed with the treatment breast milk mAbs at physiologic anti-HIV Env breast milk antibody concentrations (13). Additionally, animals were passively infused with the treatment mAbs prior to challenge to mimic placentally transferred maternal antibodies. As placentally transferred IgG would result in systemic anti-HIV titers persisting for several months postpartum, we re-infused animals 1 week after the first infusion to sustain systemic mAb titers. Of note, our use of the breast milk mAbs for the passive infusions as opposed to systemically-isolated maternal mAbs is a limitation of this model. However, HIV Env-specific mAbs isolated from milk B cells only appear distinct from those in blood in their enrichment of IgG1 isotype (14). Importantly, all animals demonstrated detectable mAb levels in saliva at 1 hour after the first mAb infusion, which was the time of the first SHIV challenge, indicating that the treatment mAbs were present at mucosal sites to mediate their potentially protective functions. Interestingly, animals from all treatment groups exhibited variable clearance of the treatment mAbs (**Figure 2**). While this finding is unlikely to alter conclusions regarding the protective capabilities of these treatments, disparities in mAb concentrations late in the study could convolute our assessments of the impact of these mAbs on post-infection SHIV viral loads in tissues at 8 weeks post challenge.

As expected, serum from groups treated with mAbs DH377 and/or DH378 exhibited neutralization against MW965 and serum from groups treated with mAb DH382 exhibited ADCC against the challenge SHIV. The lack of SHIV-1157ipd3N4 neutralization by DH378-infused animals was expected as mAb levels of DH378 did not reach the IC_50_ for SHIV-1157ipd3N4 neutralization (79µg/mL), given the limitation of the infusion volume in infant RMs. Several viremic animals treated with the control mAb developed natural neutralization responses against MW965 by week 8, and an ADCC response against the challenge SHIV in 2 of 8 animals. Yet, viremic animals from both anti-HIV mAb-treated groups failed to exhibit adaptive ADCC responses at week 8. This lack of development of ADCC functionality in serum at week 8 post challenge in the anti-HIV mAb-treated animals could indicate mAb blocking of immunogenic epitopes, but further exploration is warranted to better understand this result.

Plasma viremia was assessed to classify the extent to which anti-HIV breast milk mAb-treated animals were protected from SHIV acquisition relative to the control mAb CH65-infused animals. Importantly, as maternal mAbs employed in this study did not coevolve with the challenge virus, which would be the case in the natural setting of HIV-1 infection, the observed protection facilitated by these mAbs may underrepresent that which would occur against a truly autologous viral strain. Both the DH378-treatment and the tri-mAb cocktail treatment groups had a lower number of viremic animals compared to the control mAb-infused group (6/8 animals viremic), with only 3/6 and 2/6 animals becoming viremic over the 8 week study, respectively (**Figure 3**). Yet, this potential partial protection mediated by breast milk mAbs lacked statistical significance when compared to controls, likely due to low animal numbers. Interestingly, peak VLs, but not set point VLs, in DH378-treated, viremic animals were significantly higher than that of control animals and this difference was not attributable to more protective MHC-types in the control animals (**Table S1**). Perhaps this effect on plasma VL was due to selection of more fit variants or antibody-enhanced viral infection of target cells in CD4-blocking mAb-infused infants. However, peak VLs in the tri-mAb-treated, viremic animals were not elevated over those in control animals, indicating that a cocktail of functionally diverse mAbs may negate the potential effects of a single mAb on viral replication. Additionally, tri-mAb-treated animal LK81 demonstrated the lowest peak and set point viral loads observed in any viremic animal (**Figure 3**). This enhanced viral control post-acquisition may be attributable to the animal MHC background with 2 protective MHC alleles, Mamu-A001 and Mamu-B047a, that are both associated with lower set point viral loads (33–40). However, other viremic animals sharing the Mamu-A001 allele, including LK72 with 2 protective alleles, did not demonstrate notably lower viral loads (**Table S1**). Furthermore, these MHC alleles are not associated with decreased peak viral loads (33, 39).

Tissue-associated virus was characterized in viremic animals at 8 weeks via multiple approaches to fully characterize the state of the tissue-associated virus load within lymphoid and GI tract tissues (**Figure 4**). The size of the latent reservoir in these infant RMs was not directly assessed due to limitations of cell numbers and the lack of ART treatment prior to necropsy. Interestingly, while limited differences in measures of virus-infected cells were observed between most infant RMs regardless of treatment groups, tri-mAb-treated animal LK81 did not have detectable proviral loads or tissue mononuclear cell-associated infectious viral titers, and demonstrated low levels of active viral transcription in only 2 of 5 tissues tested. This finding is consistent with the low plasma viremia observed in this animal, and may either be due to the protective MHC alleles, the tri-mAb anti-HIV functionalities, or a combination of these factors. While our characterizations of tissue-associated virus were otherwise largely similar between treatment groups, tissue mononuclear cell-associated infectious viral titers indicated that the tonsillar tissue had heightened infectious SHIV levels compared to other lymphoid and GI tissues. This result aligns with previous reports indicating the susceptibility of tonsillar tissue as a portal of HIV entry and amplification (44–49), and further suggests that the tonsils may be a key anatomical site for strategies seeking to prevent or treat HIV infections, particularly through the oral route.

Interestingly, we observed high proviral loads in a range of tissues from several control and DH378 mAb-treated non-viremic animals (**Figure 5**). These findings may indicate either abortive or very low level SHIV infection. Generation of pseudoviruses containing 2 proviral envelope genes (*env*) isolated from these animals demonstrated that they were capable of infection in vitro, suggesting that the provirus envelope genes were functional. However, the viability of the full length provirus was not assessed due to inability to amplify the full genome. Notably, detectable provirus was not observed in CD4+ T cells isolated from any tissues of all non-viremic, tri-mAb-treated animals. This finding may indicate that the tri-mAb cocktail successfully prevented or eliminated even abortive infection in these animals. Alternatively, Liu *et al.* recently demonstrated that passive bNAb infusion mediating complete protection from intravaginal SHIV challenge in RMs did not prevent initial virus acquisition and proliferation in distal tissues, but rather that the bNAb successfully cleared this infection by around 10 days post-challenge (50). Thus, perhaps the various anti-SHIV functionalities mediated by the tri-mAb treatment in this study similarly cleared the initial infection of distal tissues. To determine whether the tri-mAb treatment prevented initial SHIV acquisition in these animals or mediated clearance of SHIV in distal tissues, additional work characterizing the viral populations present in distal lymphoid tissues at various early time points post-challenge is necessary. Regardless of the mechanism leading to uninfected distal tissues, we speculate that the ADCC-mediating mAb, DH382, may have played a pivotal role since the DH378 mAb treatment alone failed to similarly limit tissue-associated proviral loads. This speculation is supported by previous reports of an association between ADCC functionality of milk antibodies and diminished vertical transmission rates (51) and infant mortality after infection (52) in humans.

The SHIV diversity at peak viremia in infant RMs was measured to assess the capability of these maternal breast milk mAbs to limit the number of variants transmitted across the mucosa, transmitted/founders (T/F), and/or early SHIV diversification after oral challenge (**Table 1**). The fewer distinct SHIV variants isolated from tri-mAb-treated animals compared to control mAb-treated animals indicates that this cocktail of maternal breast milk mAbs successfully limited the number of T/F variants capable of establishing sustainable infection in infant RMs. Furthermore, the double protective MHC alleles in the tri-mAb-treated animals is unlikely to contribute to this trend of fewer transmitted SHIV variants, as these alleles have been associated with lower set point VLs and slowed disease progression, but not protection from virus acquisition. As DH378 mAb-treated infant RMs exhibited similar numbers of SHIV plasma variants compared to control mAb-treated animals, again the combination of distinct antibody functions is likely important for reducing SHIV variant transmission.

In conclusion, polyclonal non-broadly neutralizing maternal breast milk mAbs were capable of limiting oral virus acquisition in this infant RM model, potentially through inhibition of initial variant acquisition at mucosal sites or through rapid viral clearance in the tissues. A combination of mAbs with diverse Env binding epitopes, and neutralization and ADCC functionalities, appeared effective at lowering the number of T/F SHIV variants, and potentially SHIV levels in circulating CD4+ T cells and tissues in SHIV-challenged infant RMs. Moreover, this treatment had no association with high peak plasma VLs following breakthrough infection, as was observed with a single weakly-neutralizing mAb treatment. Additional work characterizing the state of lymphoid and GI tissues during early acute infection in the setting of non-broadly neutralizing mAbs could elucidate the exact mechanisms of protection from acquisition and/or rapid viral clearance mediated by these antibodies, which would further contribute to vaccine design approaches. While bNAbs have been repeatedly shown to be highly effective both prophylactically and therapeutically, they have also proved challenging to elicit through vaccination. Ultimately, this study confirms that even non-broadly neutralizing maternal antibodies, particularly with diverse functions, contribute to the highly inefficient infant virus transmission via breastfeeding. A better understanding of the mechanism of this protection as well as other maternal and/or infant factors that contribute to the relative protection of infants in the setting of breastfeeding may lead to more effective pediatric HIV therapeutic and prophylactic vaccine design strategies.

## Materials and Methods

More detailed description of experimental methods provided in Supplemental Methods.

### Study Animals and Specimen Collection

Twenty infant rhesus macaques (RM; 1–2 weeks old) were IV infused with either 10mg/kg anti-HIV Env gp120 monoclonal antibody (mAb) DH378 (n=6), 10mg/kg anti-influenza HA mAb CH65 (n=6), 30mg/kg α-HA mAb CH65 (n=2), or 30mg/kg of a tri-mAb cocktail composed of stoichiometric equivalents of 3 anti-HIV Env gp120 mAbs – DH377, DH378, and DH382 (n=6) - one hour prior to the first oral SHIV challenge. Animals were subjected to 3 oral challenges per day for 3 consecutive days consisting of 5,000 TCID_50_ SHIV-1157ipd3N4 (NIH AIDS Reagent Program) incubated in 1mL of RPMI containing 1µg/mL DH378, 1µg/mL CH65 (n=6), 3µg/mL CH65 (n=2), or 3µg/mL of the tri-mAb cocktail for 15 minutes, followed by dilution in ~10mL formula feed to simulate oral acquisition via breastfeeding. Animals were re-infused with their respective antibody infusions one week after the initial infusion. Blood and saliva (via weck cell sponges) were collected before each infusion, 1 hour after each infusion, 1 day after each infusion, and at weeks 2, 4, 6, and 8 of the study. All animals were necropsied at week 8 of the study and tissues were collected. Tissues were processed either fresh or after overnight shipping at 4°C to isolate mononuclear cells for sorting, or tissues were fixed in formalin for in situ hybridization. Mononuclear cells were isolated by density gradient centrifugation with Ficoll-Paque (GE Healthcare) for lymph nodes, spleen, blood, and tonsils and with Percoll (Sigma-Aldrich) for intestinal tissues, as previously described (53).

### Production of infusion mAbs

Infusion mAbs were obtained through antigen-specific B cell sorting and Ig variable gene amplification, as previously described (14), and produced through transient transfection either by the manufacturer Catalent (DH378; Catalent) or at the Duke Human Vaccine Institute in Expi293 cells (DH377, DH382, CH65), as previously described (54).

### HIV-1 Neutralization in TZM-bl Cells

Neutralizing antibody titers were measured by the reduction in Tat-regulated Luc reporter gene expression in a TZM-bl (NIH AIDS Reagent Program) reporter cell assay, as previously described (55).

### Tissue Mononuclear Cell Viral Coculture

Tissue-associated infectious virus titer was assessed through Tat-regulated Luc-F reporter gene expression to quantify infection of TZM-bl reporter cells after coculture with serial dilutions of tissue mononuclear cells isolated from RMs. For quantitative comparison, tissues were tested with a minimum of 2 replicates to employ the Reed-Meunch method to estimate the tissue-associated infectious virus titer in units of viable mononuclear cells. The cutoff for infection was defined as the maximum luminescent readout in RLU obtained from tissue mononuclear cells isolated from an uninfected animal.

### Plasma Viral RNA Load Quantification

Reverse transcriptase quantitative PCR was performed to determine the SHIV-1157ipd3N4 RM plasma RNA load, as previously described (56).

### Mononuclear Cell Provirus Quantification

RM CD4+ T cell-associated genomic DNA (gDNA) was isolated from various GI and lymphoid tissues with the QIAaMP DNA/RNA extraction kit (Qiagen) and quantified using the Biorad QX200 droplet digital PCR System according to the manufacturer instructions (Biorad) with SHIV *gag* specific primers and probe and a commercially available human TERT-specific reference assay (Biorad). The SHIV proviral load in SHIV copies/million CD4+ T cells was calculated by dividing the SHIV DNA copy number by the TERT copy number divided by 2 multiplied by 10^6^ cells.

### Measurement of virus-specific IgG levels in plasma and saliva

Enzyme-linked immunosorbent assays (ELISA) were performed as previously described (20). Plasma or saliva samples were serially diluted in 384 well ELISA plates (384 wells; Corning Life Sciences) coated with gp140 1086c, MN.gp41, Bio-V3.C, or Hemagglutinin (HA Solomon Islands; Protein Sciences Corporation). Horseradish peroxidase (HRP)-conjugated goat anti-human IgG (Jackson ImmunoResearch) was used as a secondary antibody followed by detection with SureBlue Reserve Microwell Substrate (VWR), TMB (3,3′,5,5′-tetramethylbenzidine) Stop Solution (VWR), and 450nm absorbance quantification of individual wells using a Spectramax Plus spectrophotometer (Molecular Devices). Antibody concentrations in serum and saliva were identified using purified monoclonal antibody 4-parameter standard curves of DH378, CH65, 7B2 (anti-gp41 IgG1 mAb), or the tri-antibody cocktail ranging from 0 to 300ng/mL with 3-fold dilutions.

To measure A32 blocking, plasma or saliva samples were serially diluted on 1086c gp120-coated plates prior to incubation with 200-ng/ml biotinylated A32 mAb for one hour. Inhibition of biotin-A32 binding was detected with streptavidin-HRP at 1:30,000, followed by detection as described above. A32-blocking antibody concentration in serum was determined using a 4-parameter standard curve of the tri-antibody cocktail.

### Antibody Dependent Cellular-Cytotoxicity

ADCC activity of the purified mAbs and peripheral serum samples was determined by a luciferase-based cell killing assay as previously described (31, 57). CEM.NKR_CCR5_ target cells (NIH AIDS Reagent Program) were infected with SHIV1157-ipd3N4-IMC encoding a *Renilla* luciferase reporter gene (58), followed by addition of PBMCs from a healthy HIV-seronegative donor in wells of a 96-well plate and incubation with serial dilutions of RM serum. ADCC activity (percent specific killing) was calculated from the change in RLU (ViviRen luciferase assay; Promega) resulting from the loss of intact target cells in wells containing effector cells, target cells, and serum or mAb samples compared to amounts in control wells containing target cells and effector cells alone. ADCC activity is reported as either the maximum percent killing observed for each sample or the ADCC endpoint titer, which is defined as the last dilution of serum that intersects the positive cutoff (15% specific killing) after subtraction of the average non-specific background activity observed for plasma collected prior to infusion of mAb from all available animals (n=12).

### Flow Cytometry

RM PBMCs or tissue mononuclear cells were stained with a panel of fluorochrome-conjugated antibodies and either phenotyped using an LSRII flow cytometer or bulk sorted into CD4+ and CD8+ T cells using a FACS Aria II cytometer (BD Biosciences). CD4+ and CD8+ T cells were positively selected from isolated tissue mononuclear cells by sequential selection of lymphocytes, FSC and SSC singlets, viable cells, CD45+ leukocytes, CD3+ T cells, and CD4+ vs CD8+ T cells. Data analysis was performed using FlowJo software (TreeStar).

### In Situ Hybridization

In situ hybridization for the detection and quantification of SHIV gag RNA in formalin-fixed tissue blocks was performed using the Affymetrix protocol according to manufacturer instruction (Affymetrix). Tissue blocks were stained for CD3, DAPI, and SHIV gag RNA and imaged on slides with a Leica TCS SP8 confocal microscope. The number of SHIV RNA producing cells was reported as the number of SHIV *gag* RNA+ cells per 1,000 CD3+ T cells.

### CD4+ T cell transcriptome analysis

cDNA was generated from RNA extracted from RM PBMCs and tissue mononuclear cells using the Fluidigm cDNA Prep Reverse Transcriptase Mix according to the manufacturer’s instructions and preamplified for 20 cycles. Multiplex qPCR was performed on a 48x48 Gene Expression IFC chip (Biomark) using Fast TaqMan Assays (Biomark) with the TaqMan Fast Advanced MasterMix (ThermoFisher Scientific) on the Biomark HD Instrument to quantify RNA levels of 48 genes. Raw data was uploaded into the Fluidigm Real Time PCR Analysis Software to generate Ct values. Heat map visualization was obtained using the R statistical language with the gplots visualization package.

### Transmitted/Founder Analysis

Transmitted/Founder (T/F) viral sequences in plasma were obtained by single genome amplification (SGA) through nested reverse transcriptase PCR with subsequent direct amplicon sequencing, as previously described (59). Sequence alignments and phylogenetic trees were constructed using clustalW and Highlighter plots were created using the tool at www.lanl.gov.

To identify and enumerate T/F variants, the following conditions were applied. Clusters of related sequences were visually analyzed using phylogenetic trees (Figtree v1.4) and sequences containing <2 mutations were considered a single variant. Variants containing ≥2 mutations were considered as progeny of distinct T/F genomes. Potential G-A hymermutations caused by APOBEC 3G/3F were identified using Hypermut algorithm 2.0 and were reverted for analysis if there were ≤ 2 present. Sequences that had >3 potential APOBEC 3G/3F mutations were not considered for T/F analysis (Hypermut, http://www.hiv.lanl.gov) (60). Sequence clusters of ≥ 2 sequences with ≥2 shared mutations were considered as distinct T/F variants.

### Provirus env cloning and pseudovirus preparation

SHIV *env* amplification from genomic DNA (gDNA) extracted from infant RM CD4+ T cells was done through bulk nested PCR and cloning into pcDNA3.1/V5-His-Topo (Invitrogen). Pseudovirus was produced via cotransfection of the SHIV *env* plasmid and a plasmid containing a subtype B *env* deficient HIV genome (SG3Δ*env*) in 293T cells (Invitrogen) as previously described (55). The infectivity of pseudotyped viruses was screened by single round infection of TZM-bl cells followed by detection of Tat-regulated luminescence with the Bright-Glo luciferase reagent (Promega) and infectivity was reported as RLU.

### Statistical Analysis

Statistical tests were performed with SAS v9.4 (SAS Institute). Comparisons of viral load, proviral load (copies/million cells), and the number of T/F variants in infants from each mAb treatment group were performed using the exact Wilcoxon test. The proviral load (binary designation), and the number of infants with detectable SHIV in each mAb treatment group were compared with Fisher’s exact test. False discovery rate (FDR) p-value correction was used to correct for multiple comparisons. A p-value of <0.05 (two-tailed) was considered as significant for all analyses.

## Acknowledgements

We thank Ruth Ruprect and the Dana Farber Cancer Institute for generously permitting the use of the SHIV-1157ipd3N4 for challenging the animals in this study through the NIH AIDS reagent Program. We also thank the Duke Human Vaccine Institute Protein Production Facility for help with mAb production, the Duke University Sequencing and Genomic Technology core facility for help with the ddPCR and Fluidigm assays, and David O’Connor and Roger Wiseman at the Wisconson National Primate Center for performing the MHC-typing. TZM-bl cells and SG3Δ*env* were provided by John Kappes and Xiaoyun Wu through the NIH AIDS Reagent Program. This work was funded by HHS | National Institutes of Health (R01AI1063980). The funders had no role in study design, data collection and analysis, decision to publish, or preparation of the manuscript.

## Supplemental Figure Legends

**Figure S1. Decay of passively infused mAbs in serum of tri-mAb cocktail-treated infant RMs.** Concentrations of infused mAbs in serum from pre infusion to 8 weeks post infusion are depicted for tri-mAb cocktail-treated animals. **A)** 1086C. gp140, **B)** 1086C. V3 peptide, and **C)** A32-blocking ELISAs were employed to estimate the relative concentrations of DH377 and DH382 within the tri-mAb cocktail-treated animals over the course of the study. Black arrows indicate systemic mAb infusions at days 0 and 7. ND indicates not detectable.

**Figure S2. CD4+ T cell counts in blood of mAb-treated, SHIV-1157ipd3N4 challenged infant RMs. A)** Absolute and **B)** relative CD4+ T cell counts in blood of mAb-treated and orally SHIV-challenged infant RMs collected longitudinally. Red, blue, and green symbols indicate control mAb-, DH378-mAb, and tri-mAb-treated animals, respectively.

**Figure S3. Tissue-associated infectious SHIV levels measured through tissue mononuclear cell coculture with TZM-bl reporter cells.** Mononuclear cells isolated from tissues 8 weeks after oral SHIV-1157ipd3N4-challenge of control mAb-, DH378 mAb-, and tri-mAb-treated infant RMs were serially diluted and cocultured with TZM-bl reporter cells for 72hrs, followed by luminescent detection of tissue-associated SHIV infectivity in relative luminscence units (RLU). The RLU limit of detection for positive tissue-associated SHIV infection (dashed line) was defined as 3 standard deviations above the mean maximum RLUs elicited from PBMCs of unchallenged control RMs (n=3) in the coculture assay.

**Figure S4. CD4+ T cell proportions in tissues of mAb-treated, SHIV-1157ipd3N4 challenged infant RMs. A)** Proportion of CD4+ T cells of total T cells and **B)** total CD45+ mononuclear cells in various lymphoid and GI tissues of mAb-treated and orally SHIV-challenged infant RMs at necropsy (8 weeks post infusion). Red, blue, and green symbols indicate control mAb-, DH378-mAb, and tri-mAb-treated animals, respectively. Squares indicate non-viremic animals, while circles indicate viremic animals.

**Figure S5. Summary of SHIV detection and proportion of animals with detectable cell-free or cell-associated SHIV in breast milk mAb-infused, orally SHIV-challenged infant RMs. A)** Binary indication of detectable plasma SHIV RNA, peripheral CD4+ T cell SHIV provirus, and tissue CD4+ T cell SHIV provirus detection. For each animal, SHIV detection methods with detectable SHIV in any tested tissue or at any timepoint were indicated with a red “+”, while those with undetectable values in all tested tissues and at all timepoints were indicated with a grey “-”. **B)** Proportion of animals with detectable cell-free or cell-associated SHIV as defined by detectable SHIV plasma RNA, SHIV provirus in tissues or PBMCs, tissue mononuclear cell-associated infectious SHIV titers, or SHIV RNA-transcribing cells.

**Figure S6. Heatmap hierarchical clusterings of CD4+ T cell RNA transcription profile in infant RMs.** Heatmap visualization of a Fluidigm-based multiplex RT-qPCR assay to quantify expression of a 27-gene panel selected to represent activation and cell death pathways in genomic RNA from FACS-isolated **A)** PBMC and **B)** lymph node CD4+ T cells. A two-dimensional hierarchical clustering heatmap was constructed in the R programming language. Green represents a high column Ct Z score, or low RNA quantities while red represents low Ct values, or high RNA quantities. Cell-associated proviral copies, quantitated by ddPCR as log *gag* copies/10^4^ CD4+ cells, and animal treatment group are depicted with color-coded bars to the right of the clustering heatmap. The proviral load color scale ranges are determined by the rank quartile *gag* quantities.

**Figure S7. Representative phylogenetic trees of SHIV *env* variants isolated from passively mAb-infused, SHIV-1157ipd3N4- challenged infant RMs.** Standard SGA techniques were used to isolate the SHIV envelope gene from plasma of **A)** control mAb-treated, **B)** DH378 mAb-treated, and **C)** tri-mAb-treated infant RMs, and amplicons were sequenced. Clustalw and the Kimura 2 Parameter (K2P) method were used to align amplicons and construct phylogenetic trees rooted at the SHIV-1157ipd3N4 envelope gene (Acc. No. DQ779174), respectively. Distinct variant clusters (V; indicated by bold lines) within each animal were defined as having ≥2 unique mutations that were observed in ≥2 amplicons.

**Figure S8. Phylogenetic tree analysis of SHIV *env* isolated from passively breastmilk mAb-infused infant RMs 2 weeks after oral SHIV challenge.** Standard SGA techniques were used to isolate SHIV *env* genes from each animal 2 weeks following oral SHIV challenge and amplicons were sequenced. Clustalw and Kimura 2 Parameter (K2P) method were used to align amplicons and construct the phylogenetic tree rooted at the SHIV-1157ipd3N4 stock *env*, respectively. Amplicons from control mAb-treated, DH378 mAb-treated and tri-mAb-treated animals are colored red, blue, and green, respectively. The scale bar represents 0.0008 nucleotide mutations per site. The black arrow indicates the SHIV-1157ipd3N4 challenge *env* sequence. SHIV proviral *env* sequences isolated from CD4+ T cells in LH19 and LG73 submandibular LNs at 8 weeks post infection were included in the tree and are indicated by *.

